# Description of *Venenivibrio orakeikorakoensis* sp. nov. and *Venenivibrio aotearoaensis* sp. nov, and emended description of genus *Venenivibrio*, *Venenivibrio stagnispumantis* species

**DOI:** 10.64898/2026.09.07.749006

**Authors:** Holly E. Welford, R. Carlo Carere, Meghan E.A. Marshall, Kirill Lagutin, Kevin A. Mitchell, Matthew B. Stott

**Affiliations:** Te Kura Pūtaiao Koiora | School of Biological Sciences, Te Whare Wānanga o Waitaha | University of Canterbury, Ōtautahi | Christchurch, Aotearoa | New Zealand; Te Tari Pūhanga Tukanga Matū | Department of Chemical and Process Engineering, Te Whare Wānanga o Waitaha | University of Canterbury, Ōtautahi | Christchurch, Aotearoa | New Zealand; Callaghan Innovation, Te Awa Kairangi ki Tai | Lower Hutt, Aotearoa | New Zealand

**Keywords:** *Venenivibrio*, *Aquificota*, New Zealand, geothermal, hot spring

## Abstract

Two thermophilic bacterial strains, designated OKO1 and KUI1, were isolated from hot springs within the Taupō Volcanic Zone of New Zealand. Analysis of the 16S rRNA gene identified both as members of the genus *Venenivibrio*, sharing 99.13% and 98.82% sequence similarity with *Venenivibrio stagnispumantis* CP.B2^T^, respectively. The novel isolates were characterised using a polyphasic taxonomic approach alongside a recharacterization of *V. stagnispumantis*. Despite their close 16S rRNA gene sequence similarity, the strains differed markedly in cell morphology, physiology and genomic content. Both new isolates exhibited broad tolerance to temperature and pH, while re-examination *V. stagnispumantis* CP.B2^T^ revealed substantially wider physiological capabilities than previously reported, bringing its phenotype into closer agreement with the new isolates. Comparative genomic analyses identified key differences in genes encoding cytochromes, hydrogenases, and nitrogen metabolism, suggesting distinct metabolic potential among the strains. Although the genomes shared average nucleotide identity (ANI) values of 95-96%, placing them near the accepted species boundary, the combined phenotypic and genomic evidence support recognition of OKO1^T^ and KUI1^T^ as two species within the genus. We therefore propose *Venenivibrio orakeikorakoensis* sp. nov. (type strain OKO1^T^, ACCESSION) and *Venenivibrio aotearoaensis* sp. nov. (type strain KUI1^T^, ACCESSION) as new members within the genus, and emend the descriptions of *Venenivibrio stagnispumantis* and the genus *Venenivibrio* to incorporate the expanded phenotypic and genomic diversity revealed in this study.

## Introduction

The phylum *Aquificota* encompasses a diverse range of species found generally in extreme environments globally, including geothermal springs and marine hydrothermal vents (Reysenbach, 2015, Gupta, 2014). The phylum presently includes ∼32 known species (Freese, *et al*., 2026) across three families, *Aquificaceae*, *Desulfurobacteriaceae* and *Hydrogenothermaceae*, all of which are thermophilic, with growth ranges varying between 38.5 °C and 95 °C (Power, *et al*., 2024, Deckert, *et al*., 1998). Most species within *Aquificota* utilize the Knallgas (2H_2_(g)+ O_2_(g)→2H_2_O(l) + energy) reaction for energy generation (Gupta, 2014, Gupta and Lali, 2013), with a few capable of heterotrophic growth on sugars or other organic compounds (Takai, *et al*., 2001, Caldwell, *et al*., 2010, Nakagawa, *et al*., 2005). Many species also utilize sulfur compounds in metabolic reactions (Reysenbach, 2015, Gupta, 2014, Cao, *et al*., 2017, McKay, *et al*., 2022). Members of the phylum *Aquificota* contribute to several key biogeochemical processes, including carbon cycling through carbon dioxide fixation (Hügler, *et al*., 2007), nitrogen fixation (Palmer, *et al*., 2025), and sulfur cycling (McKay, *et al*., 2022). Carbon dioxide (CO_2_) is fixed via two differing citrate cleavage methods within the reductive TCA (rTCA) cycle (Hügler, *et al*., 2007), with the members of *Aquificaceae* employing citryl-CoA synthetase and citryl-CoA lyase (E.C 4.1.3.34) whereas both *Hydrogenothermaceae* and *Desulfurobacteriaceae* utilizing ATP citrate lyase (E.C 2.3.3.8). Of the three families within *Aquificota*, all *Desulfurobacteriaceae* are reported as obligate anaerobes using reduced sulfur compounds or nitrate as electron acceptors. Conversely members of the families *Aquificaceae* and *Hydrogenothermaceae* are capable of utilizing oxygen, nitrate, ferric iron, or reduced sulfur compounds as terminal electron acceptors (Reysenbach, 2015, Gupta, 2014).

Within the family *Hydrogenothermaceae*, cultivated representatives are typically microaerophilic, and utilize hydrogen, reduced sulfur compounds electron donors and, in some cases, nitrate as acceptors for energy conservation (Götz, *et al*., 2002, Stöhr, *et al*., 2001, Hetzer, *et al*., 2008, Takai, *et al*., 2003). The family *Hydrogenothermaceae* currently comprises four formally described genera: *Persephonella*, *Sulfurihydrogenibium*, *Hydrogenothermus,* and *Venenivibrio*. All characterised *Hydrogenothermus*, *Persephonella*, and *Venenivibrio* species utilise H_2_ as an electron donor, as well as three of the five published *Sulfurihydrogenibium* species (Nakagawa, *et al*., 2005, Götz, *et al*., 2002, Hetzer, *et al*., 2008, Takai, *et al*., 2003, Stohr, *et al*., 2001, Aguiar, *et al*., 2004, Flores, *et al*., 2008, O’Neill, *et al*., 2008, François, *et al*., 2021, Nakagawa, *et al*., 2003).

Representatives of the genera *Hydrogenothermus* and *Persephonella* have only been detected within marine-based ecosystems (Ferrera, *et al*., 2014, Mino, *et al*., 2013) with all representatives isolated from hydrothermal vents and chimneys. Conversely, to date, both *Venenivibrio* and *Sulfurihydrogenibium* have only been detected in terrestrial geothermal systems (McKay, *et al*., 2022, Power, *et al*., 2024, Hedlund, *et al*., 2015, Hug, *et al*., 2014). The sole representative of the genus *Venenivibrio*, *Venenivibrio stagnispumantis* CP.B2^T^, was first isolated from Champagne Pool, Waiotapu, Aotearoa-New Zealand (Hetzer, *et al*., 2008). Hetzer *et al*. (2008) reported it to be a hydrogenotrophic microaerophile with an obligate requirement for sulfur or thiosulfate. It was reported to grow across a pH and temperature range of 4.8-5.8 and 45-75 °C, with growth optima of pH 5.4 and 70 °C. However, in a recent study by Power *et al*. (2024), the authors postulated the genus *Venenivibrio* was likely endemic to Aotearoa-New Zealand and, on the basis of occurrence in physicochemically diverse hot springs, proposed that the type strain’s temperature and pH growth range were broader than previously reported. In addition, the authors reported up to 110 *Venenivibrio* OTUs in a near national survey of Aotearoa-New Zealand hot springs.

Here, we describe two novel species within the genus *Venenivibrio* (OKO^T^ and KUI1^T^), both isolated from hot springs in the Taupō Volcanic Zone (TVZ) of Aotearoa-New Zealand, for which we propose the names *Venenivibrio orakeikorakoensis* sp. nov. and *Venenivibrio aotearoaensis* sp. nov. respectively. We also recharacterized *V. stagnispumantis*, both expanding and clarifying its phenotypic and genomic characteristics. Consequently, we emend the descriptions of both the genus *Venenivibrio* and the species *V. stagnispumantis* to reflect the broader diversity revealed in this study.

## Materials and methods

### Sampling and isolation

Unless otherwise stated, mMSH medium (Hetzer et al., 2008) was used to cultivate and test *Venenivibrio* strains. This medium consists of (per L): KCl; 0.50 g, MgCl_2_×6H_2_O; 1.36 g, MgSO_4_×7H_2_O; 7.00 g, NaS_2_O_3_×5H_2_O; 2.00 g, CaCl_2_×2H_2_O; 0.40 g, NH_4_Cl; 0.20 g, KH_2_PO_4_; 0.25 g, MES; 1.95 g, sulfur; 0.25 g, and 10 mL trace mineral solution (containing Na-EDTA×2H_2_O; 500 mg, CoCl_2_×6H_2_O; 150 mg, MnCl_2_×4H_2_O; 100 mg, FeSO_4_×7H_2_O; 100 mg, ZnSO_4_; 118.5 mg, KAl(SO_4_)_2_; 142.2 mg, Na_2_WO_4_×2H_2_O; 30 mg, CuSO_4_×5H_2_O; 32.24 mg, Ni_2_SO_4_×6H_2_O; 20 mg, H_3_BO_3_; 20.7 mg, H_2_SeO_3_; 7.46 mg, Na_2_MoO_4_×2H_2_O; 10 mg. The medium was adjusted to pH 5.5 using 1 M NaOH. All media were prepared in 120 mL borosilicate serum vials sealed with a butyl rubber bung (50 mL medium: 70 mL headspace). The medium was sterilized via autoclave at 105 °C (15 p.s.i, for 15 minutes) to avoid elemental sulfur melting and coalescence. After autoclaving, nitrogen gas was sparged through the medium for one minute per 10 mL of medium, then a H_2_:CO_2_:O_2_ gas mix were added via 0.22 µm filtered gastight syringe in the ratios of 79:16:5 without replacement of nitrogen for a final constitution of 50:39.5:8:2.5 (v/v) of N_2_:H_2_:CO_2_:O_2._ Cultures were incubated at 70 °C without shaking unless otherwise stated.

Data from the 1000 Springs Project database (Power, *et al*., 2018) were used to identify hot springs within the TVZ that exhibited the greatest diversity and relative abundance of *Venenivibrio* OTUs. These sites were subsequently targeted for the enrichment and isolation of novel *Venenivibrio* strains. Two springs were selected, Kuirau Park feature 87, and Orakei Korako feature 9. *In situ* pH, conductivity, and temperature were measured using an EcoSense pH100A meter (YSI, OH, USA), an EcoSense EC300A meter (YSI, OH, USA), and a Fluke 51 II thermocouple (Fluke, WA, USA), respectively. Geographic coordinates were recorded using a Garmin GPS 60 receiver (Garmin, KS, USA). Water and sediment samples were collected in sterile 500 mL Nalgene bottles from Kuirau Park (176° 14’ 33.27087”S, 38° 07’ 54.58016”E, pH 6.26, 66.2 °C, 2182 µS cm⁻¹) and Orakei Korako (176° 08’ 32.06024”S, 38° 28’ 29.82073”E, pH 6.72, 79.8°C, 2520 µS cm⁻¹) (Figure S1) and stored at 4 °C until required. To enrich for *Venenivibrio* strains, a 10% (v/v) inoculum of the water–sediment slurry was added to 45 mL of sterile mMSH medium in 120 mL serum vials using a sterile syringe. Cultures were incubated at 70 °C for 7 days without agitation. Growth was monitored by phase-contrast microscopy (Zeiss Primostar).

Enrichments were then serially diluted to extinction into borosilicate Balch tubes (10 mL volumes) three times until an axenic culture was achieved. Two isolates were obtained via this process, *Venenivibrio* sp. OKO1^T^ and *Venenivibrio* sp. KUI1^T^ from Orakei Korako and Kuirau Park respectively. Culture purity was determined using phase-contrast microscopy and 16S rRNA gene sequencing.

### DNA extraction, PCR, and sequencing

The putative taxonomic identities of strains OKO1^T^ and KUI1^T^ were determined by sequencing their 16S rRNA genes. In addition, the 16S rRNA gene of the type strain, *Venenivibrio stagnispumantis* CP.B2^T^, was re-sequenced to confirm its identity and resolve ambiguous nucleotide positions present in the original reference sequence (DQ989208.1). Actively growing cultures were grown to stationary phase, with cell pellets collected via centrifugation (11,000 × g, 10 min) and suspended in 800 µL of Qiagen DNeasy Powersoil CD1 buffer. DNA was then extracted using Qiagen DNeasy PowerSoil kit as per the manufacturer’s protocol (Qiagen, Hilden, Germany), except for the cell lysis step, which was performed using a horizontal vortex adapter. PCR was performed using the universal primers 9F (5’ GAGTTTGATCMTGGCTCAG 3’) and 1492R (5’ ACGGYTACCTTGTTACGACTT 3’) targeting the 16S rRNA gene (thermocycler Eppendorf Mastercycler x50), and the product cleaned using the Zymo DNA Clean & Concentrator-5 kit (Zymo Research, California, USA). Sequencing of the 16S rRNA genes were carried out via Sanger sequencing at the University of Canterbury. A similar methodology was applied for whole genome DNA extraction, except extracted gDNA was cleaned using the Zymo DNA Clean & Concentrator-25 kit (Zymo Research, California, USA) as per the manufacturer’s protocols. Whole genome sequencing was undertaken on the Illumina NovaSeq6000 SP platform generating 150 bp paired-end sequencing products by the Australian Centre for Ecogenomics (ACE). The genomes were assembled using Unicycler v.0.4.8 (Wick, *et al*., 2017) and annotated using RASTk v1.073 (Aziz, *et al*., 2008) within the KBase wrapper (Arkin, 2018). Sequence and assembly QC were assessed via QUAST v4.4 (Gurevich, *et al*., 2013). Genomes were uploaded to NCBI (accession numbers JBSPYY000000000 and JBSPYY000000000), and the 16S rRNA gene sequence of *V. stagnispumantis* CP.B2^T^ was updated (DQ989208.2; 12/5/2026)

### Taxa identification

Pairwise analysis of the *Venenivibrio* 16S rRNA gene sequences was performed via BLAST using the rRNA/ITS databases and the MegaBLAST algorithm (Altschul, *et al*., 1990). A 16S rRNA gene-based phylogenetic assessment was undertaken within the ARB ecosystem and a phylogenetic tree was constructed using TREE-PUZZLE (Schmidt, *et al*., 2002), a quartet-puzzling maximum-likelihood algorithm (10,000 puzzling steps and HKY substitution model). GTDB-tk (Chaumeil, *et al*., 2019) was used to classify the *Venenivibrio* genomes based upon 49 conserved genes generating a relative evolutionary divergence (RED) score and average amino acid identities (AAI) against the type strains of the genera of *Hydrogenothermaceae*. Average nucleotide identities (ANI) were generated using FastANI v0.1.3 (Jain, *et al*., 2018). The phylogenomic tree was created using FastTree 2 (Price, *et al*., 2010). Genes were identified through word searches and investigating RAST subsystems (Aziz, *et al*., 2008, Brettin, *et al*., 2015). Predicted hydrogenase amino acid sequences were classified using the HydDB (v1.0) web server (Søndergaard, *et al*., 2016).

### Phenotypic characterization

Unless otherwise stated, strains KUI1^T^, OKO1^T^, and *V. stagnispumantis* CP.B2^T^ were phenotypically characterised in triplicate and growth was determined via phase contrast microscopy. Optical density measurements were not used due to rapid floc formation and the presence of elemental sulphur in standard mMSH growth medium.

Temperature growth ranges were determined using a Terratec temperature gradient test tube oscillator from 38.5 °C to 81.5 °C with temperature increments of ∼1.25 °C (2 × 24 tubes in total; experiments performed in duplicate). Inoculated Balch tubes were incubated for 14 days and were visually inspected daily for growth. pH tolerances were determined by inoculating each strain into mMSH media ranging from pH 3.0-8.5 in 0.5 increments. Media were prepared and gassed as previously described with the omission of MES buffer, except for between pH 5.5-6.5. A sodium citrate/citric acid buffer was used to maintain pH between 3.0-5.0, and HEPES buffer was used to maintain pH between 7.0-8.5. pH was adjusted using either 0.1 M HCl or 0.1 M NaOH. Tubes were incubated for one week and growth was checked daily. The final pH of each culture was confirmed using Universal Indicator pH indicator paper (Macherey-Nagel, Düren, Germany). Growth across a range of salinities (0-10% w/v; NaCl) was tested in mMSH medium in concentrations of 0%, 0.5%, 1%, 2.5%, 5%, and 10% w/v over a 14 day period. For each of the observed maximum and minimum pH, temperature, and salinity ranges for each strain, isolates were sequentially subcultured (3×) into the standard mMSH medium and grown at the standard pH, temperature and salinity conditions (70 °C, pH 5.5, 0% NaCl) to confirm viability of the tested condition.

Gram stain, catalase, and oxidase tests were carried out using standard protocols. Enzymatic activity of each strain was tested using API**®** ZYM test strips (bioMérieux product 25200, Marcy-l’Étoile, France) via the following method: Late exponential phase *Venenivibrio* cultures were centrifuged at 11,000 × g for 5 minutes to form a cell pellet. The pellet was then resuspended in 1.5 mL of UltraPure water (Invitrogen product 10977015, MA, USA), and then approximately 65 μL of this culture suspension was added to each well. The test strips were then incubated at 37 °C and color changes observed after 4.5 hours incubation. *Escherichia coli* CMB44 was used as a control.

For each species, the potential usage of various carbon, nitrogen and sulphur sources were determined via the following methods. For carbon-sources, CO_2_-free mMSH medium was used with the addition of (0.5 g/L; 0.22 μm syringe filter sterilised) glucose, xylose, maltose, sucrose, cellulose, chitin, sodium formate, disodium succinate, sodium bicarbonate, or casamino acids. CO_2_ was omitted from the headspace by flushing each tube with N_2_ for one minute per 10 mL of medium. H_2_ and O_2_ were added for a final headspace constitution of 50% N_2_: 47.5% H_2_: 2.5% O_2_ (v/v). Each tube that displayed growth was sequentially subcultured three times to confirm the result. For nitrogen sources, Balch tubes containing NH_4_Cl-free mMSH medium were made anoxic through headspace vacuuming (five minutes), followed by flushing the headspace with argon gas to displace any trace N_2_. H_2_:CO_2_ was added in the ratio of 80:20, followed by 5% O_2_ (v/v) for a final ratio of 50:39.5:8:2.5 (v/v) of Ar:H_2_:CO_2_:O_2_. Alternative sources of nitrogen were added in the following concentrations: sodium nitrate; 1 mM, urea; 30 mM, casamino acids; 7.4 mM, sodium nitrite; 1 mM, proline; 10 mM, or ammonium chloride; 3.7 mM. Prior to inoculation, 10 mL cultures were centrifuged at 8,000 × g for ten minutes. Nine mL of the supernatant was removed and replaced with N-free medium to dilute residual ammonium chloride. Each culture that displayed growth was subcultured three times in the same medium to ensure growth was not due to carryover of NH_4_Cl. To test alternate sulfur-sources, mMSH medium was made with the omission of MgSO_4_×7H_2_O, Na_2_S_2_O_3_, and elemental sulfur. Alternative sulfur compounds were added in concentrations as follows; elemental sulfur; 3% w/v (936 mM), Na_2_S_2_O_3_×5H_2_O; 40 mM, Na_2_S×9H_2_O ; 2.1 mM, L-cysteine hydrochloride; 1.9 mM, methionine; 3.4 mM, cystine; 2.1 mM, pyrite; 0.5g/10 mL (w/v), Na_2_S_4_O_6_×2H_2_O; 2.3 mM, polysulfide; 0.4 mM, KSCN; 10 mM, C_2_H_3_NaO_2_S; 0.9 mM, Na_2_SO_3_; 10 mM. Prior to inoculation, 10 mL cultures were centrifuged at 2,000 × g for one minute to remove elemental sulfur, then centrifuged at 8,000 × g. for 10 minutes. Nine mL of the supernatant was removed from the cell pellet and replaced with sulfur-free medium. This was then used as an inoculum into media containing the sources of sulfur. Tubes that displayed growth were sequentially subcultured three times to confirm viability.

For each strain, H_2_ (47.5% v/v), elemental sulfur (3 g/L), and thiosulfate (2 g/L) were tested as sole electron donors, each in the absence of the other donor. To determine the oxygen growth requirements of the *Venenivibrio* isolates, Balch tubes containing mMSH medium with 0.002 g/L (w/v) resazurin were prepared by flushing with N_2_ gas until resazurin colour change was observed (pink to clear). Anaerobically prepared L-cysteine hydrochloride (3.2 mM) was then added as a reducing agent to remove traces of oxygen. Headspaces were then injected with H_2_/CO_2_, with O_2_ in concentrations of 0%, 1.25%, 2.5%, 3.75%, 5%, 7%, 10%, and 20% (v/v). Media were inoculated with 10% v/v inoculum and incubated for one week. Cultures displaying positive growth at the upper and lower O_2_ concentrations were subcultured three times at these conditions to confirm results.

Dissimilatory sulfate reduction was determined using Hach reagents (SulfaVer4) and analyzed using a Hach colourimeter (DR900; Hach Company, Colorado, USA). mMSH medium was prepared as previously described omitting thiosulfate or elemental sulfur, MgSO_4_×7H_2_O was reduced to 3.5 g/L in congruence with literature on general sulfate reducers concentrations (e.g. (Bagheri Novair, *et al*., 2024)). The medium was rendered anaerobic by supplementation of ascorbic acid (0.01%, v/v) as a reducing agent to scavenge residual oxygen, with the resazurin (0.002 g L⁻¹) used as an oxygen indicator. H_2_:CO_2_ were added to the headspace as previously described. Cultures were inoculated and incubated at 70 °C for 7 days. The contents of one SulfaVer4 powder pillow was added to each tube as per the manufacturer’s instructions. Each was then diluted into sterile medium 100-fold to measure within the detection-limits and analysed. Dissimilatory nitrate reduction was determined using HACH reagents (NitraVer5) with a HACH colourimeter (DR900) (Hach Company, Colorado, USA). This was undertaken using a cadmium reduction test for the presence of nitrate, in comparison to a non-inoculated negative control. Cultures were prepared as previously described with no sulfur, thiosulfate, or oxygen present and a decreased MgSO_4_ concentration (3.5 g/L). Instead, 1 mM of sodium nitrate was added as a terminal electron acceptor. Resazurin (0.002 g/L) and ascorbic acid (0.01% v/v) were added as described above. H_2_/CO_2_ were added as described previously. Media were inoculated using 10% v/v liquid culture of each isolate, then incubated for 7 days at 70 °C. After 7 days, the contents of a Hach NitraVer5 Powder Pillow were added to each tube as per manufacturer’s instructions and results were analysed using the Hach colourimeter (DR900) (Hach Company, Colorado, USA). Sample values were subtracted from the controls to determine the amount of nitrate consumed. Phase-contrast microscopy was also used to determine any change in cell numbers.

For imaging cells using transmission electron microscopy, cultures of strains KUI1^T^ and OKO1^T^, and *V. stagnispumantis* CP.B2^T^ (40 mL) were centrifuged at 5,000 × g for 2 minutes. Pellets were resuspended in 1 mL of Karnovsky’s fixative, then centrifuged again at 5,000 × g. for 5 minutes. Pellets were resuspended in 0.1 M sodium cacodylate buffer for 5 minutes, 3 times. Samples were then suspended in 3% (w/v) ultra-low melting point agarose and centrifuged for 5 minutes at 5,000 × g., then cooled at 4 °C. The samples were then fixed using 1% w/v osmium tetroxide in 0.1 M phosphate buffer for 2 hours before being washed in 0.1 M phosphate buffer then ultrapure water three times each for 5 minutes. They underwent a graded degradation from 25-95% (v/v) for 15 minutes at each concentration. Epoxy resin was then added to the samples and cured for 22 hours at 60 °C. Ultrathin sections (100 nm) were cut using an EM UC7 ultramicrotome then collected on 200-mesh copper grids, which were then stained using 2% (w/v) uranyl acetate for 5 minutes, then followed with 0.2% (w/v) lead citrate for 45 seconds. A Talos 120C transmission electron microscope (Thermofisher Scientific, USA) was used to visualise the prepared grid samples. Negative staining was carried out by coating Formvar 200-mesh copper grids with a carbon layer of 3 nm and charged with a glow discharge treatment for 10 seconds. A 10 μL aliquot was pipetted on the grid which was then washed with water and dried with filter paper. The grid was placed face down onto 8 μL of 1% (w/v) uranyl acetate for 10-30 seconds. Grids were then dried for 15 minutes and visualised using a Talos 120C transmission electron microscope (Thermofisher Scientific, USA).

### Fatty acids and quinones analysis of *Venenivibrio* strains

To prepare cell biomass for quinone and fatty acids analyses, *V. stagnispumantis* CP.B2^T^ was batch cultivated in mMSH medium (pH 5.5) within a Labfors 5 Microbial Bioreactor at a working volume of 2.0 L. Following sterilization of the autoclave (121 °C, 15 p.s.i., 60 minutes), cultures were incubated at 70 °C with mixing (500 r.p.m.) and 200 mbar overpressure. To minimize evaporative liquid loss, inlet gases were humidified via passage through 500 mL MilliQ water prior to filter sterilisation and exited via a water-cooled gas condenser. Throughout cultivation, the concentration of dissolved O_2_ was maintained at 2% (v/v) saturation via variable airflow within a headspace otherwise balanced by H_2_/CO_2_/N_2_ (4:20:76, v/v) supplied at 25 mL/min via mass flow control. Stationary phase cultures were harvested and stored at 4 °C before being processed. To concentrate the biomass, 1 L aliquots were centrifuged at 15,000 × g for 30 minutes at 4 °C. The resulting cell pellets were then transferred into a 50 mL centrifuge tube and frozen at -20 °C. *Venenivibrio* strains OKO1^T^ and KUI1^T^, cultures were grown in 1 L Schott bottles with butyl rubber bungs with 400 mL of mMSH medium as described previously and the same headspace ratios as described above for OKO1^T^ and KUI1^T^ Cultures were grown for up to three weeks, then cells were collected via filtration using a 0.22 µm vacuum filter.

Fatty acid methyl esters for all three strains were prepared from biomass according to the protocol detailed by (Svetashev, *et al*., 1995). Fatty acids were analyzed using an Agilent 7890 gas chromatograph equipped with flame ionization detector and DB-wax UI column (30 m × 0.25 mm i.d., 0.25 µm). Helium was used as the carrier gas, 1.1 mL/min; the split ratio was 1:30. Injector and detector temperatures were 25 °C and oven temperature was 205 °C. Fatty acids were identified by equivalent chain length (ECL) values (Stránský, *et al*., 1992), and comparison with standards.

Respiratory quinones were extracted from the *V. stagnispumantis* CP.B2^T^ biomass using a modified version of the method described by Nishijima et al. (1997). Five to 30 mg of freeze-dried cell material was ground to a fine powder using a mortar and pestle, extracted by addition of 2:1 chloroform/methanol followed by ultrasonication (20 °C, 15 min). The extract was filtered through a 0.2 µm nylon syringe filter and dried under a stream of nitrogen. Analysis of extracts were carried out using a modified version of the method described by Nishijima et al. (1997). Extracts were dissolved in acetone (ca. 200 µL) and analysed by Liquid Chromatography-Mass Spectrometry (LCMS). Low resolution targeted liquid chromatography tandem mass spectrometry (LC-MS/MS) was carried out on a system comprising of a Shimadzu 8040 LCMS system equipped with photodiode array detector and triple quadrupole low resolution MS detector utilizing an Atmospheric Pressure Chemical Ionization (APCI) source for ionisation. High resolution accurate mass LCMS was carried out on a Waters Premier G2XS QTof instrument with APCI source. The same chromatographic conditions were used for both high and low resolution LCMS analysis. Chromatography was carried out on a Waters Acquity BEH C18 1.7-µm 2 × 150 mm column at 30 °C. A linear gradient elution profile was employed, initial elution conditions were 100% methanol which was held for 1 min, the percentage of 2-propanol was then increased to 5% at 4 min from sample injection, then increased linearly to 100% at 22 min, and held at this concentration for 1 min before being reduced to initial conditions over 1 min. The column was equilibrated for 7 min prior to a subsequent injection being made. A flow rate of 0.2 mL/min was employed. Respiratory quinones were identified in the UV-Vis and MS TIC chromatograms using a combination of their characteristic UV-Vis online absorption spectra, their characteristic MS/MS fragmentation pattern, and molecular formula derived from accurate mass analysis of the observed ions (expected *m*/*z* within 5 ppm of predicted).

## Results and discussion

### Enrichment, isolation, and identification

Novel *Venenivibrio* isolates were obtained from samples collected from hot springs at Kuirau Park and Orakei Korako, designated KUI1^T^ and OKO1^T^, respectively (Figure S1). Strain KUI1^T^ exhibited a slightly curved rod morphology with characteristic run-and-tumble motility. Phase-contrast microscopy (Figure 2a–c) showed cells measuring 3.0–4.5 µm in length and 0.8–1.4 µm in width. In contrast, strain OKO1^T^ consisted of substantially larger, straight rod-shaped cells measuring 5.0–5.9 µm in length and 0.9–1.1 µm in width and displayed a characteristic spinning motility. TEM analysis showed multiple flagella present for each strain (Figures S3-S5). The genus was named for their vibrio shaped-cells, however of the three strains, only the type strain appears to possess this vibrio-like morphology. Transmission electron microscopy (Figure 1, Figures S2–S5) revealed that both strains possessed multiple flagella and a typical diderm cell envelope comprising inner and outer membranes. While only a qualitative observation, KUI1^T^ cultures produced a pronounced sulfide odor, whereas cultures of CP.B2^T^ and OKO1^T^ had no discernible odor.

**Figure 1:**
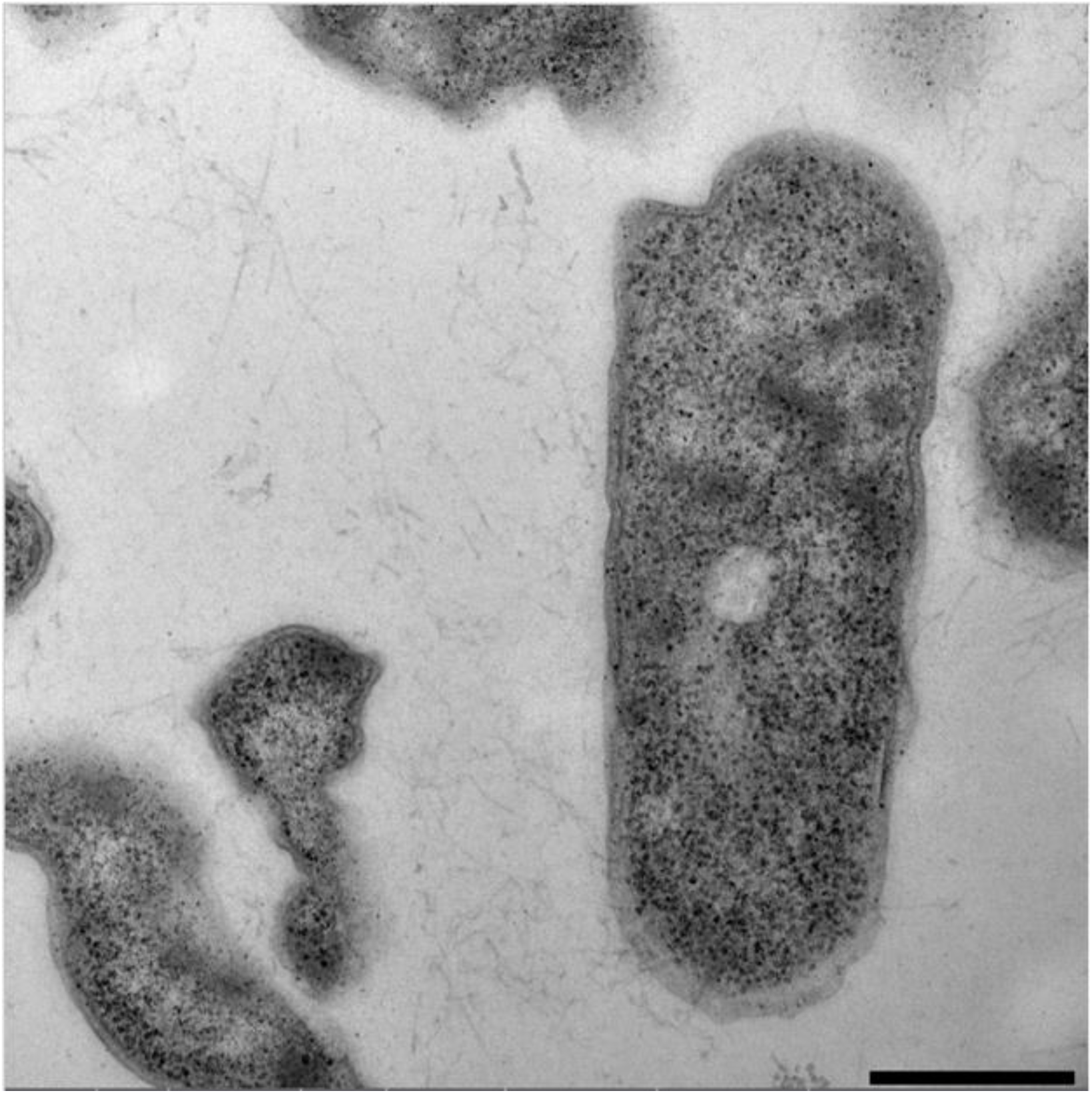
Transmission electron micrograph of *Venenivibrio orakeikorakoensis* OKO1^T^ thin section (0.5% w/v uranyl acetate, 10 sec exposure time) at 22,000 × magnification. Scale bar represents scale of 500 nm.

**Figure 2:**
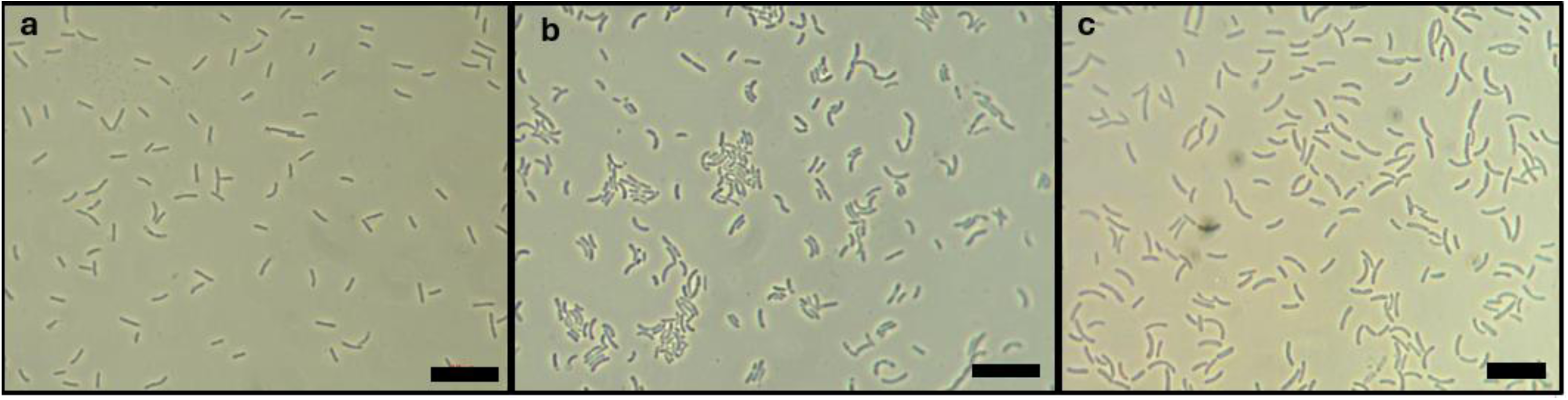
Phase contrast microscopy images of (a) strain KUI1^T^, (b) *V. stagnispumantis* CP.B2^T^, and (c) *Venenivibrio* strain OKO1^T^. Scale bar 10 µm. Images taken at 1000 × oil immersion.

Pairwise comparisons of the full-length 16S rRNA gene sequences strains KUI1^T^ and OKO1^T^ gave sequence similarity scores of 98.82% and 99.13% to *V. stagnispumantis* CP.B2^T^ (NR_044029.1) respectively, with the next closest sequences being that of a New Zealand hot spring (Kuirau Park) clone (99.26% and 99.13%; AF402979) and *Sulfurihydrogenibium azorense* Az-Fu 1^T^ (94.64% and 94.65%; NR_102858.1) (Table S1). Maximum-likelihood phylogenetic analysis placed both isolates within the genus *Venenivibrio* which formed a well-supported clade between the *Hydrogenothermaceae* genera, *Persephonella* and *Sulfurihydrogenibium* (Figure 3). To resolve discrepancies observed during sequence alignment, the 16S rRNA gene of the type strain, *V. stagnispumantis* CP.B2^T^, was also re-sequenced. Comparison with the published sequence identified eight previously unresolved nucleotide positions. The cited sequence similarity scores and phylogenetic placement uses the updated 16S rRNA gene sequence (DQ989208.2; updated on 12/5/2026).

**Figure 3:**
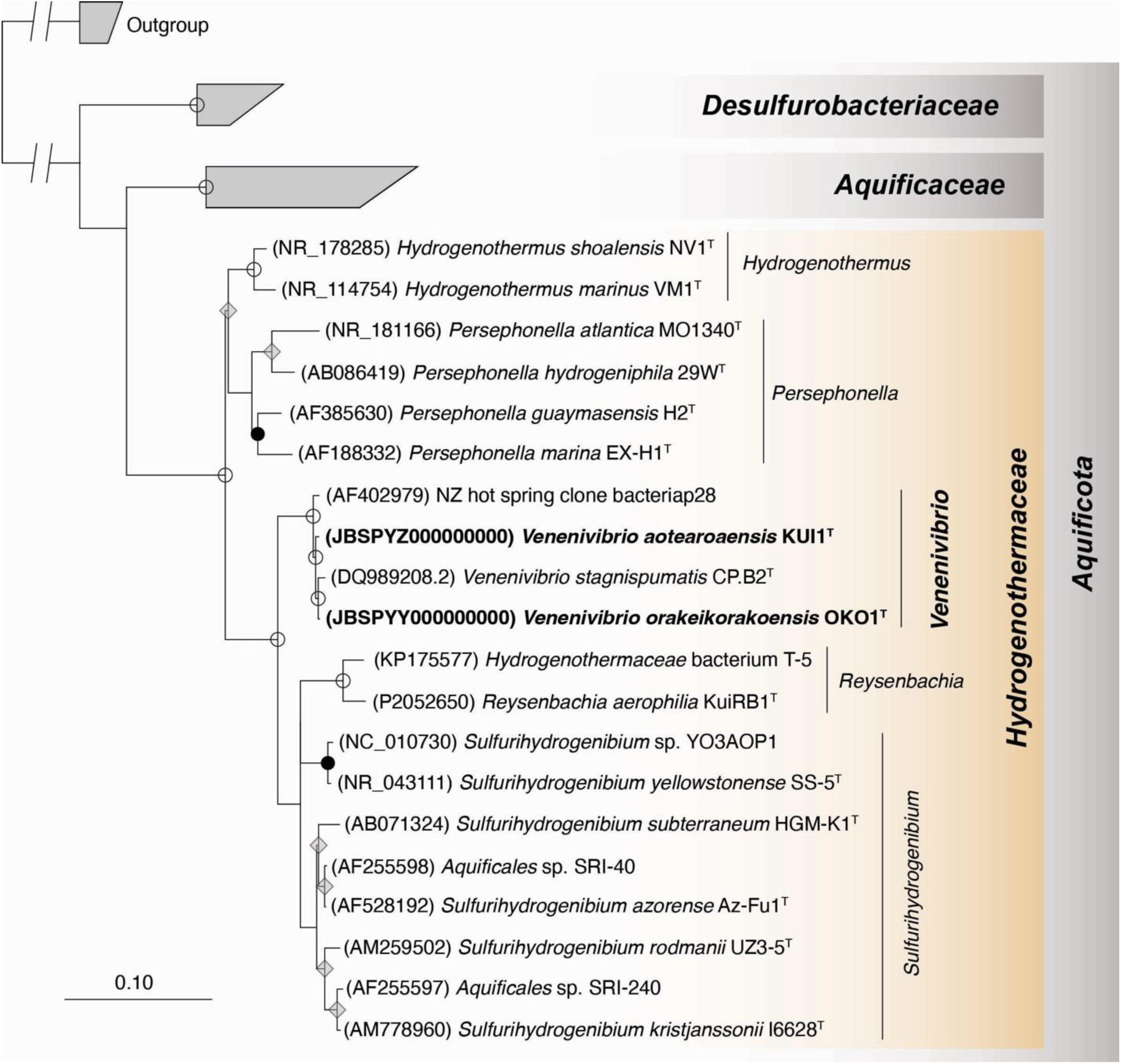
Maximum-Likelihood16S rRNA gene-based phylogenetic tree created using TREE-PUZZLE, a quartet-puzzling maximum-likelihood algorithm (10,000 puzzling steps and HKY substitution model)., within the ARB ecosystem. This tree demonstrates the phylogenetic relationship between the *Venenivibrio* strains discussed in this study (CP.B2^T^, OKO1^T^, and KUI1^T^). The outgroup of this tree were three members of the *Methanocaldococcus* genus, *Methanocaldococcus indicus* SL43^T^ (AF547621), *Methanocaldococcus jannaschii* JCM10045 (AB603516), *and Methanocaldococcus lauensis* SG7^T^ (MK602249). A preprint is available for *Reysenbachia aerophila* KUI-RB^T^ (PZ052650) (Marshall, *et al*., 2026).

### Whole Genome analysis

The genomes of the two new *Venenivibrio* isolates were sequenced and annotated, resulting in genomes of 1,690,398 bp, and 1,735,438 bp for KUI1^T^ and OKO1^T^ respectively (Table S2). Both genomes were larger than that of the type strain (1,566,079 bp (Power, *et al*., 2024, Power, *et al*., 2023)) and had %G+C contents between 29.15-29.2 mol%, a value consistent with taxa within the family *Hydrogenothermaceae* (GCF_003688665.1; (Gupta, 2014, Stohr, *et al*., 2001). Whole genome sequence comparison of the type strain with strains KUI1^T^ and OKO1^T^ using the ANI metric gave values of 95.35% and 95.30% respectively, and 96.74% between KUI1^T^ and OKO1^T^. These results indicate that all three taxa sit within the genus *Venenivibrio* and have genome dissimilarities great enough to warrant the delineation of three novel species. The *Venenivibrio* genomes clustered in a monophylogenetic clade distinct from neighbouring genera, *Sulfurihydrogenibium, Persephonella* and *Hydrogenothermus* (Figure 4) and was consistent with the 16S rRNA gene phylogenetic placement (Figure 3). Whole genome comparison via the AAI metric of the three *Venenivibrio* representatives with the type strains of other *Hydrogenothermaceae* genera resulted in values of between 76.92-77.2% to *S. azorense*, the closest phylogenetic relative, further confirming previous conclusions that genus *Venenivibrio* is phylogenetically distinct from that of genus *Sulfurihydrogenibium*.

**Figure 4:**
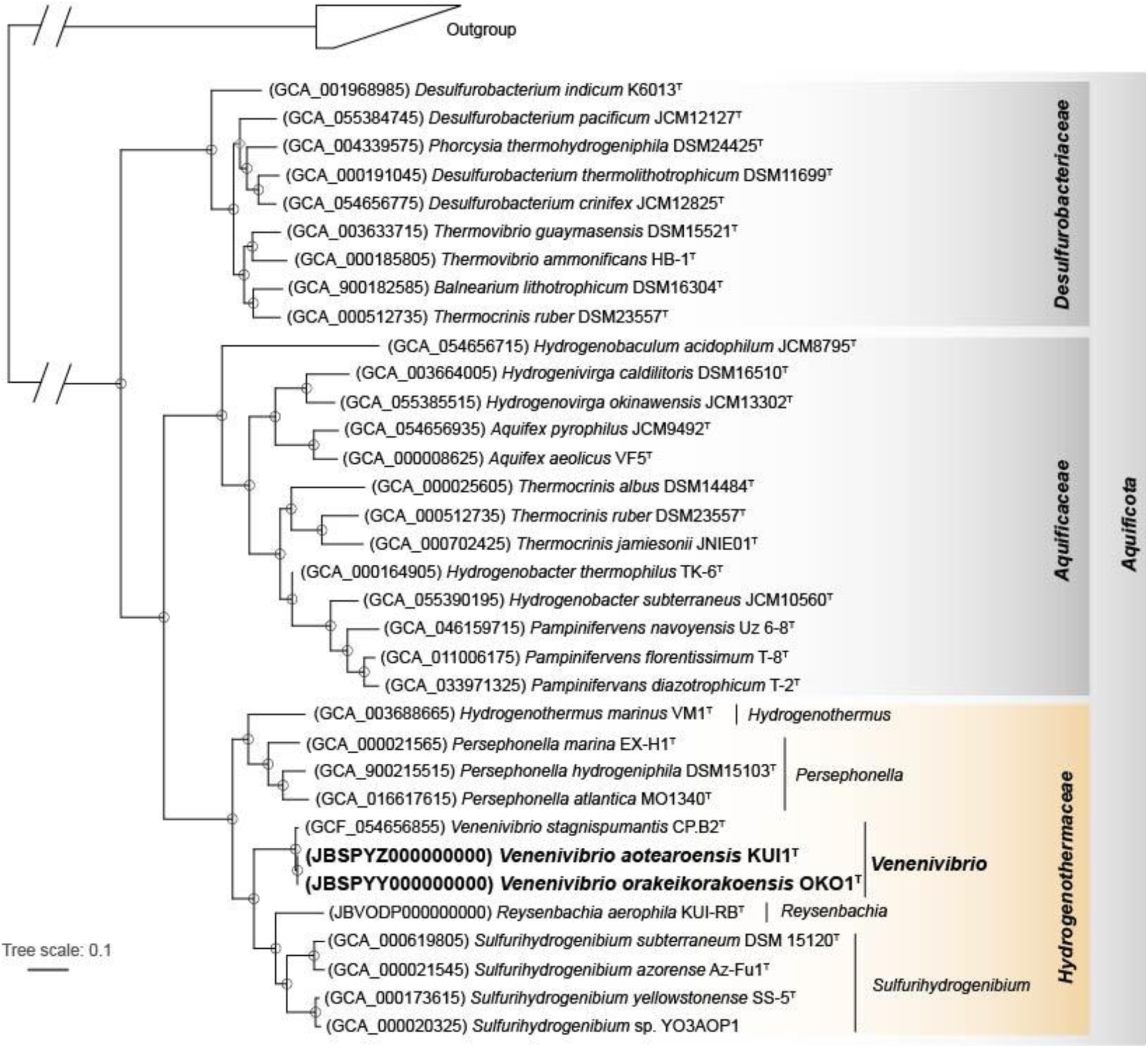
Whole genome maximum-likelihood phylogenomic tree displaying the relationship between members within the phylum *Aquificota*. The tree was generated using FastTree 2 based on the alignment of genomes (Price et al., 2010). Representatives of *Methanocaldococcus* were used as an outgroup (*M. jannaschii* DSM 2661 GCF_000091665, *M. lauensis* SG7 GCF_902827225, and *M. indicus* SL43 GCF_049554935). A preprint for *Reysenbachia aerophila* KUI-RB^T^ is available (Marshall, *et al*., 2026).

Analysis of 16S rRNA gene sequence similarity and whole-genome ANI places the *Venenivibrio* strains examined in this study at or near the accepted genomic threshold for species delineation (Beye, *et al*., 2018, Hackmann, 2025, Jain, *et al*., 2017, Ciufo, *et al*., 2018). Despite their close genomic relatedness, comparative genome analyses revealed substantial differences in genes associated with energy metabolism, including hydrogen oxidation, sulfur metabolism, nitrogen cycling, and terminal respiration. These differences indicate that relatively small marker gene nucleotide divergence can correspond to marked variation in metabolic potential, suggesting that closely related *Venenivibrio* species have diversified to exploit distinct ecological niches within geothermal environments. Using numerical thresholds, particularly that of the 16S rRNA gene similarity, as a hard limit for species differentiation has been debated (Murray, *et al*., 2021). Instead, it is suggested that species demarcation is not a universal threshold, due to variations in horizontally acquired genes and associated divergence events (Palmer, *et al*., 2019). In this regard, genetic diversity can be considered a continuum, and identifying novel species requires a polyphasic approach that includes the consideration of genomic, phylogenetic, metabolic and phenotypic characteristics (Varghese, *et al*., 2015).

The metabolic potentials of the three strains were investigated by the screening of genomes for key respiration genes. The hydrogenotrophic capability was confirmed with multiple respiration hydrogenases identified in each genome. All three genomes encoded the *Aquificota*-specific cytosolic NiFe H₂-uptake hydrogenases (Group 2d), which are suspected to play a role in supporting carbon fixation, but the broad functional role is currently unknown. In addition, strain KUI1^T^ encoded a membrane-bound H₂-uptake hydrogenase associated with anaerobic nitrate and sulfur respiration (Group 1b), while strain OKO1^T^ encoded a bidirectional cytosolic hydrogenase that may participate in sulfur reduction (Group 3b) (Greening, *et al*., 2016).

The compliment of terminal oxidase cytochromes present in the genomes of each strain also differed among the three strains, suggesting variation in terminal electron acceptors and respiratory strategies across oxygen gradients. The high affinity of bd-type oxidases for O₂ were encoded in the genomes of all three strains, presumably enabling growth under microaerophilic conditions (Popovic, *et al*., 2010, Borisov, *et al*., 2011) and was consistent with the growth characteristics observed for all three strains in this study. As members of the *Aquificota* commonly occur at high abundance in geothermal biofilms and streamers (Tan, *et al*., 2025), these respiratory adaptations likely facilitate persistence across the steep oxygen gradients characteristic of such environments. Both OKO1^T^ and KUI1^T^ genomes also encoded nitrate reductases and transporters for both assimilatory and dissimilatory nitrate reduction, which are both absent in the *V. stagnispumantis* CP.B2^T^ genome (Palmer, *et al*., 2025, Esclapez, *et al*., 2014). This suggests OKO1^T^ and KUI1^T^ are likely to possess the ability to utilise a range nitrogen species as nitrogen sources and also as part of anaerobic respiration (Pajares and Bohannan, 2016). Dissimilatory sulfate pathway genes were not identified in any of the genomes sequenced. Interestingly, both KUI1^T^ and OKO1^T^ encode sulfur oxidation pathways that may allow for sulfur oxidation or assimilation. The presence of genes encoding the sulfhydrogenase enzyme complex for all three strains suggests that sulfur reduction may be coupled to hydrogen oxidation (Greening, *et al*., 2016, Ma, *et al*., 1993, Kanai, *et al*., 2011). The presence of this complex may explain the limited anaerobic growth using sulfur as a terminal electron acceptor observed in the characterisation experiments of KUI1^T^ and OKO1^T^ (see below) and the sulfide odour observed in KUI1^T^ cultures.

Finally, the OKO1^T^ genome encodes genes for a complete Mo-Fe nitrogenase enzyme (*nifHDK*), with the full regulatory operon (*nifABENXU*, *ntrYC*, and sensors) for nitrogen fixation (Esclapez, *et al*., 2014, Pajares and Bohannan, 2016, González, *et al*., 2006, Chan and Wheatcroft, 1993, Zumft, 2005). Genes for nitrogen fixation were not present in either KUI1^T^ or *V. stagnispumantis* CP.B2^T^ genomes.

### Physiology and growth substrates

Previous research into the diversity of microbial communities in Aotearoa-New Zealand geothermal ecosystems found genus *Venenivibrio* OTUs were detected in elevated abundances across a broad range of temperature and pH conditions (Power, *et al*., 2024, Power, *et al*., 2018) which were inconsistent with the grow limit parameters previously reported for the type strain (pH 4.8-5.8, pH_opt_ 5.5 and 45-75°C, T_opt_ 70 °C; Hetzer *et al*., 2008). To address these inconsistencies, we recharacterized *V. stagnispumantis* CP.B2^T^ alongside the phenotypic characterization of the two new *Venenivibrio* isolates. Both strains KUI1^T^ and OKO1^T^ displayed similar growth ranges and optima to those observed for *V. stagnispumantis* (KUI1^T^: pH_range_ 3.5-8.0, pH_opt_ 5.5-8.0, OKO^T^: pH_range_ 3.5-8.0, pH_opt_ 6.0-6.5), although strain KUI1^T^’s optimal growth pH was broader (Table 2). Substantial morphological variability was observed across this pH range for *V. stagnispumantis* CP.B2^T^, with small cocci cells present at pH 3, then single, paired, and chains of vibrio were visualised around the optimum, and finally aggregates of cells were observed around pH 7 and above. Similar morphological changes in response to agitation versus static cultivation have previously been reported for the related *Hydrogenothermaceae* species, *Persephonella hydrogeniphila* (Nakagawa, *et al*., 2003). Conversely, no morphological changes were observed in strains OKO1^T^ and KUI1^T^ under the culture conditions examined in this study. The temperature range and optima for *V. stagnispumantis* CP.B2^T^ more closely reflected its previous characterization (T_range_ 38.5-80 °C and T_opt_ 70.4 °C). As with the pH growth observations, both KUI1^T^ and OKO1^T^ had comparable temperature ranges and optima to that of the type strain (KUI1^T^; T_range_ 38.5-79.7 °C, T_opt_ 65.7 °C, OKO^T^; T_range_ 40-77.4 °C, T_opt_ 60.1-72.8 °C). All strains could tolerate a large range of NaCl concentrations, with *V. stagnispumantis* growing between 0-8% (optimum 0-0.2) (Power, *et al*., 2024), KUI1^T^ growing between 0-5% (optimum 0-0.5), and OKO1^T^ growing between 0-10% (optimum 0-1) (w/v). The NaCl tolerance of *V. stagnispumantis* CP.B2^T^ was almost 10-fold greater than what was originally reported (Hetzer, *et al*., 2008). Both the temperature, but in particular, the pH growth ranges observed of the new isolates in this study were substantially broader than were originally reported of *V. stagnispumantis* CP.B2^T^, and can partially explain the broad pH and temperature distribution in which *Venenivibrio* OTUs reported in the 1000 Spring Project survey (Power, *et al*., 2024, Power, *et al*., 2018). Furthermore, the pH and temperature optima of the three isolates match the environmental conditions where the greatest abundances of *Venenivibrio* taxa were detected (temperatures ∼37-80°C pH ∼4-7; Power et al., 2024).

**Table 1:** Fatty acid composition of *Venenivibrio stagnispumantis* CP.B2^T^, *Venenivibrio* strains OKO1^T^, and KUI1^T^. “n.d.” represents not detected. Data for *P. marina* EX-H1^T^ and *H. marinus* VM1^T^ retrieved from Götz, *et al*., (2002) and Stohr, *et al*., (2001) respectively.

|  | CP.B2<sup>T</sup> | OKO1<sup>T</sup> | KUI1<sup>T</sup> | *P. marina* | *H. marinus* |
| --- | --- | --- | --- | --- | --- |
| C<sub>12:0</sub> | 2.9 | 0.5 | n.d. | n.d. | n.d. |
| C<sub>16:0</sub> | 1.6 | 7.1 | 2.7 | n.d. | n.d. |
| C<sub>18:0</sub> | 31.1 | 56.9 | 32.6 | 16.3 | 22-24 |
| C<sub>18:1n-9</sub> | 10.5 | 5.5 | 11.0 | 16 | 15-16 |
| C<sub>18:1n-7</sub> | 1.5 | 1.6 | 2.2 | n.d. | n.d. |
| C<sub>19:0</sub> | 0.3 | 2.7 | 1.9 | n.d. | n.d. |
| C<sub>20:0</sub> | 4.2 | 6.1 | 3.9 | 6.1 | n.d. |
| C<sub>20:1n-9</sub> | 46.9 | 17.9 | 44.8 | 21 | 46-51 |
| C<sub>20:1n-7</sub> | 1.1 | 1.8 | 0.9 | n.d. | n.d. |

**Table 2:**
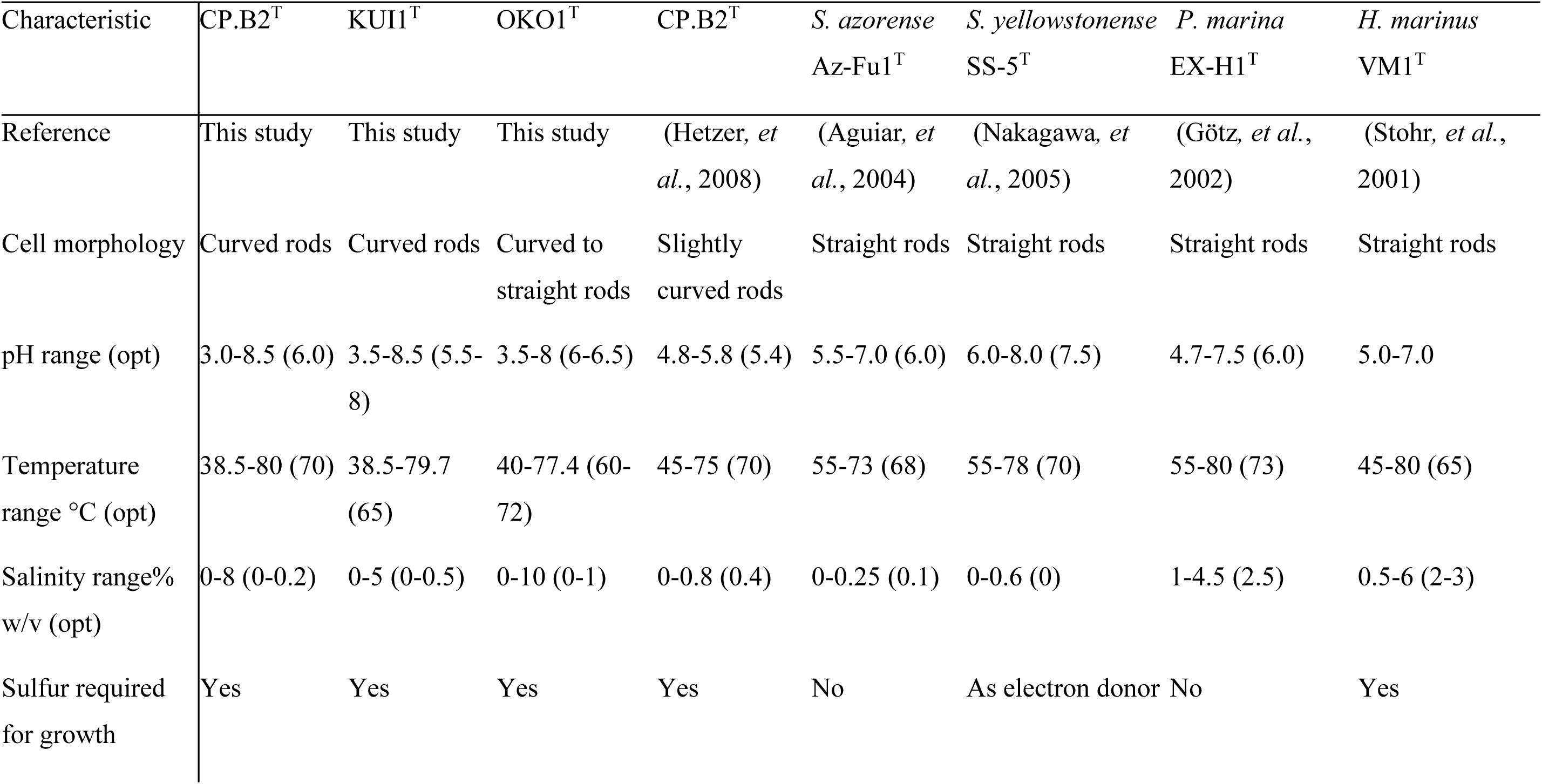
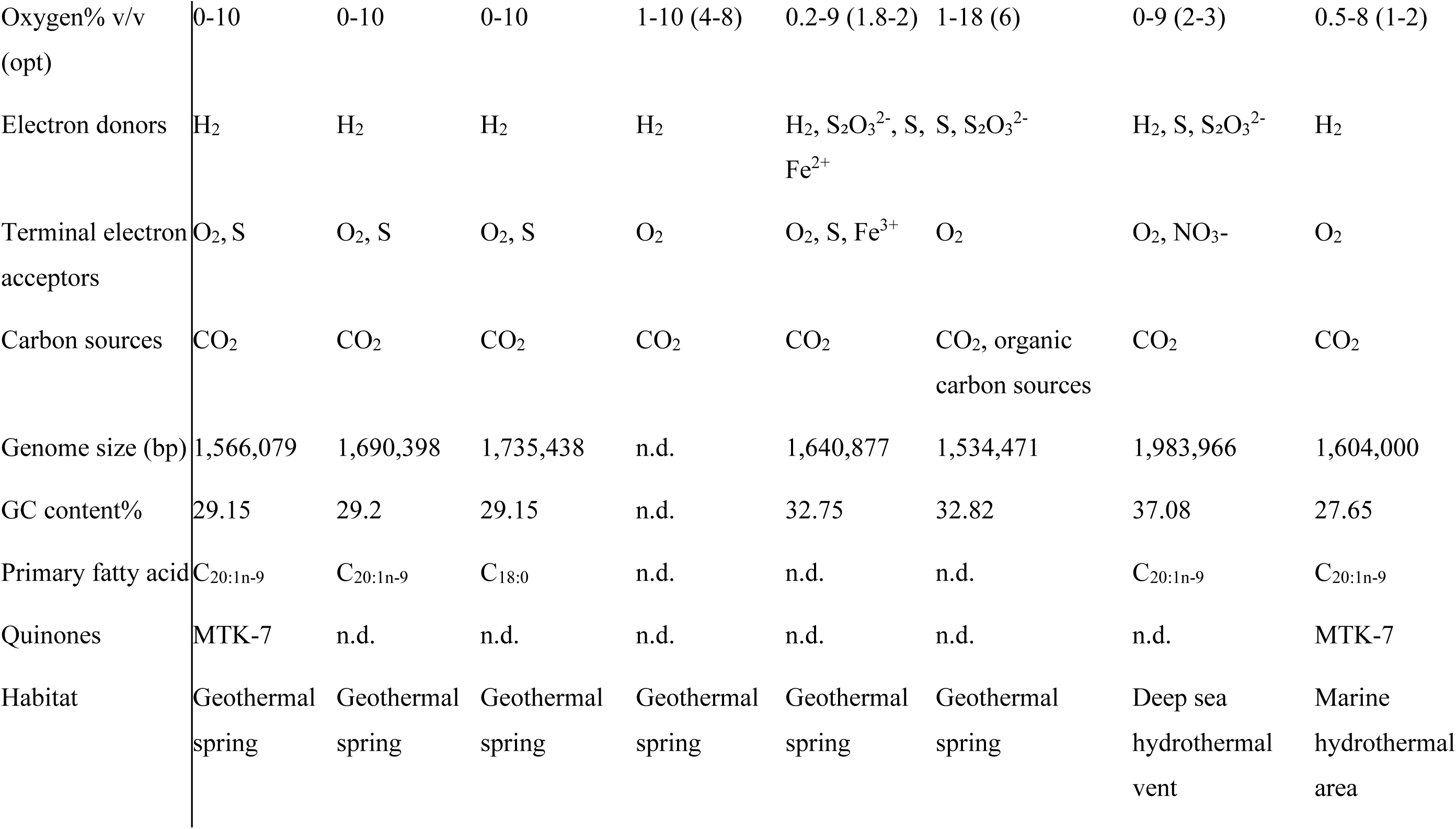
Phenotypic and genomic comparisons of type strains within the family *Hydrogenothermaceae* and novel *Venenivibrio* strains isolated from this study. The original description of *V. stagnispumantis* CP.B2^T^ (Hetzer *et al*., 2008) has also been included as a comparison. *S. azorense* data obtained from Aguiar, *et al*., (2004), *S. yellowstonense* data obtained from Nakagawa, *et al*., (2005), *P. marina* data obtained from Götz, *et al*., (2002), and *H. marinus* data obtained from Stohr, *et al*., (2001). n.d.: not determined.

**Table 3:**
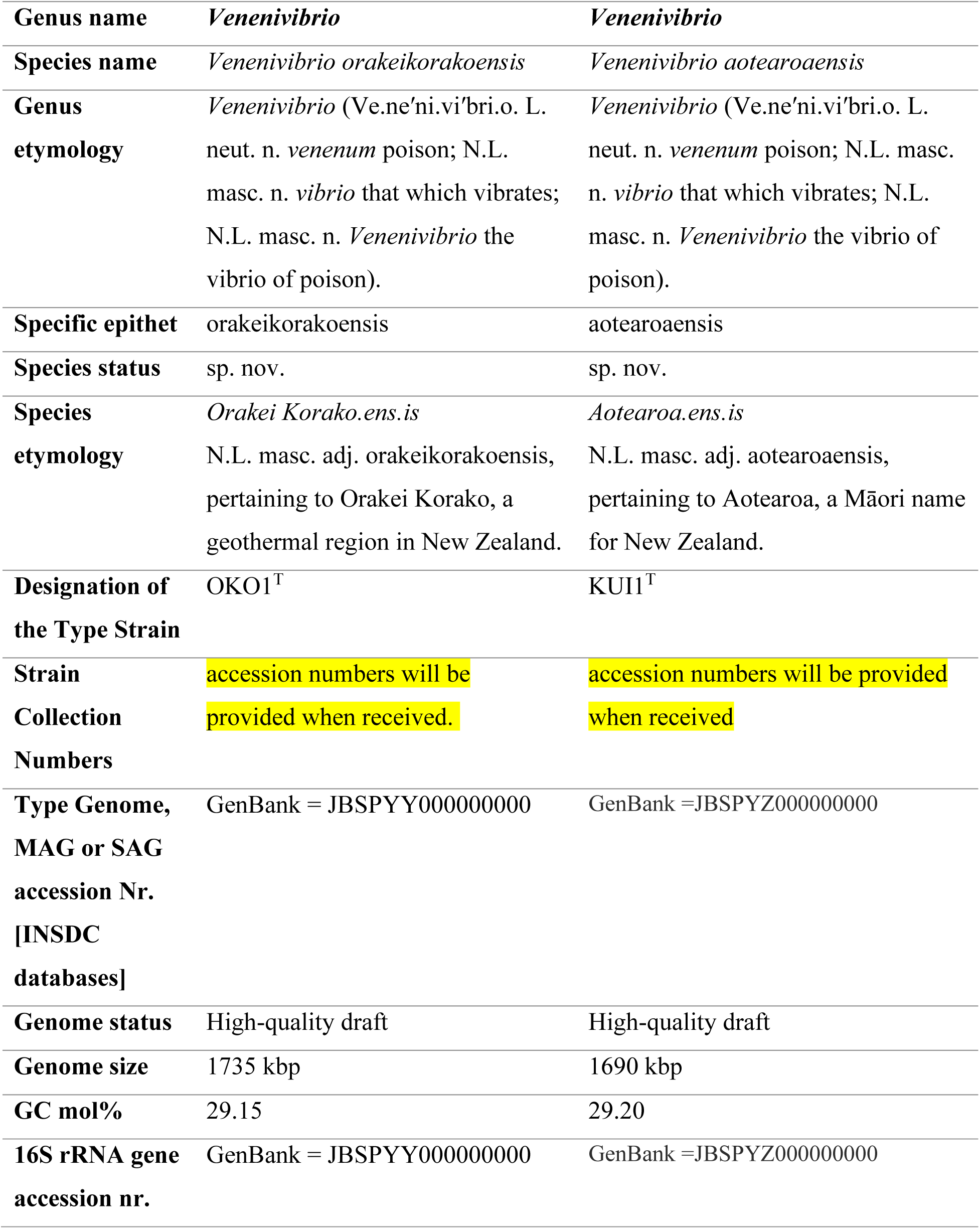
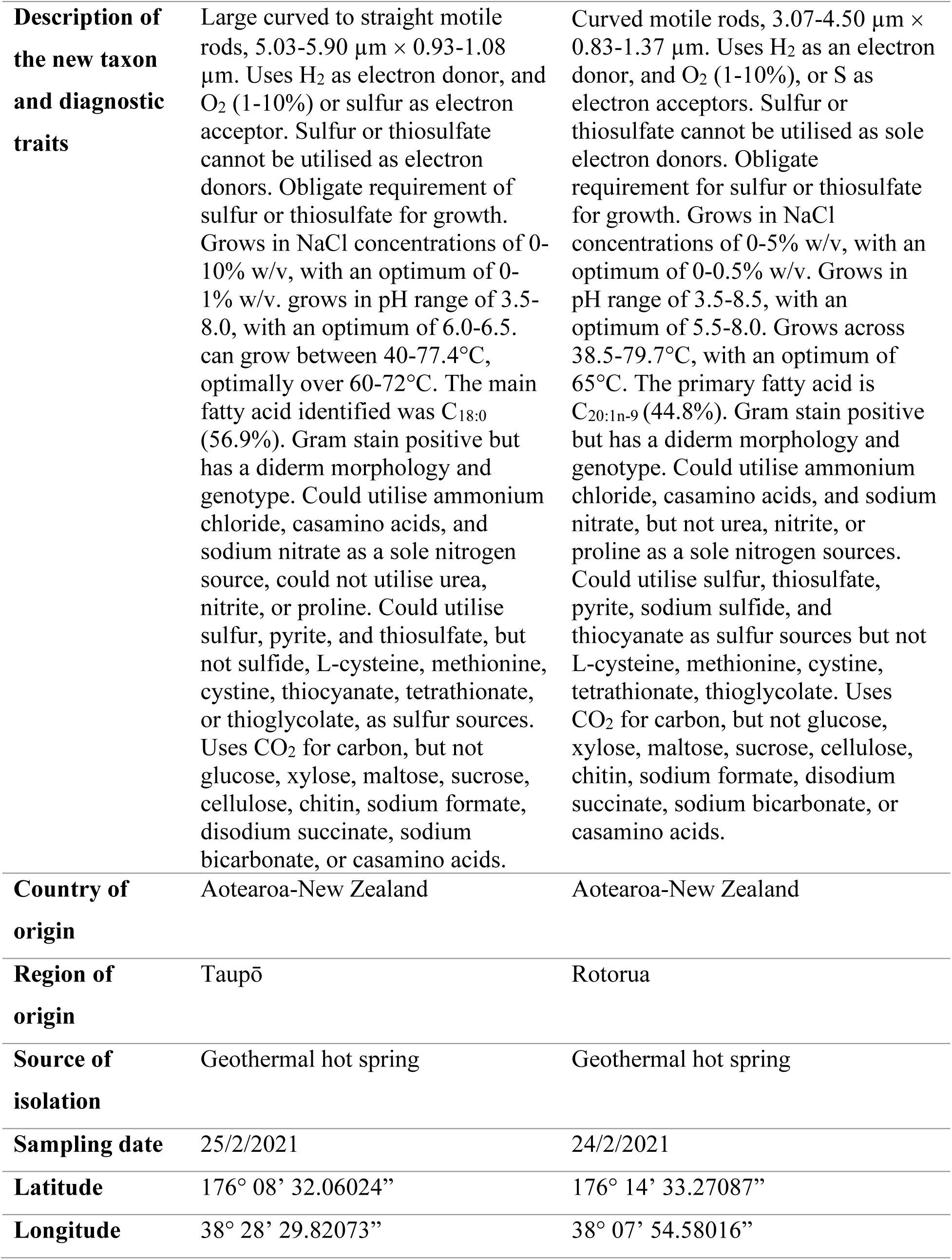
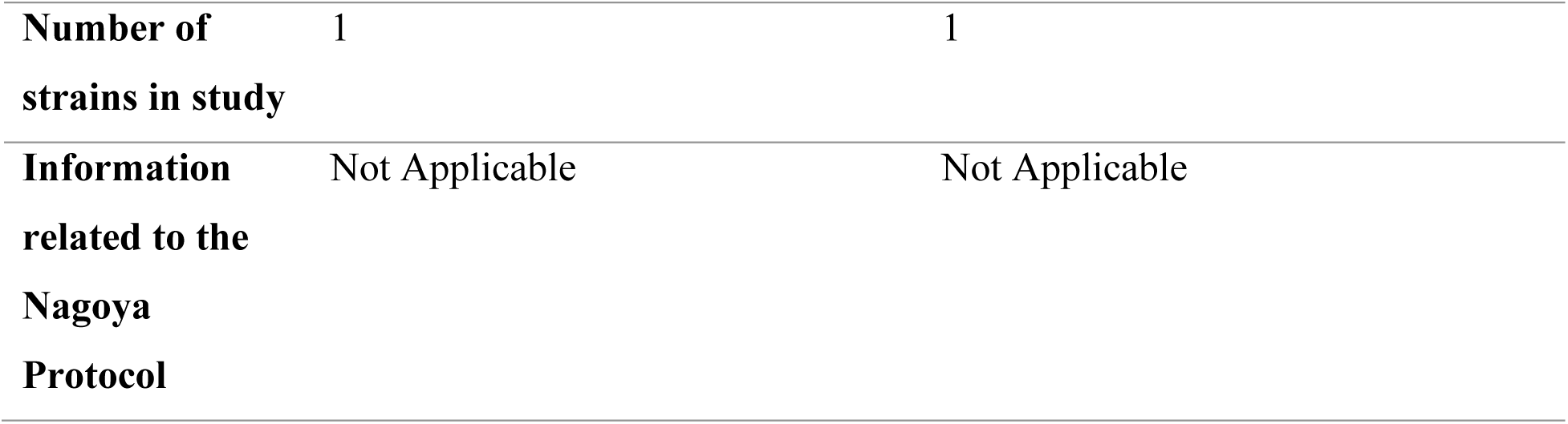
Novel species descriptions of Venenivibrio orakeikorakoensis OKO1^T^ and Venenivibrio aotearoaensis KUI1^T^.

Hydrogen gas serves as the only energy source for all strains, with no growth observed with any reduced sulfur compounds or organic substrates. All strains required CO_2_ as their source of carbon; no other carbon sources were utilised by any of the strains. CP.B2^T^ was only able to use ammonium chloride as its source of nitrogen with no growth occurring using any other sources of nitrogen. Both OKO1^T^ and KUI1^T^ grew utilising casamino acids, sodium nitrate, and ammonium chloride as sole nitrogen sources. Sulfur or thiosulfate was required for growth of all strains when using H_2_ as an energy source. CP.B2^T^ was able to utilise elemental sulfur, thiosulfate, cystine, and polysulfide as its required sulfur source, whereas, OKO1^T^ could utilise elemental sulfur, pyrite, and thiosulfate, and KUI1^T^ utilized elemental sulfur, thiosulfate, pyrite, Na_2_S, and thiocyanate.

All strains grew aerobically in microaerophilic conditions (1-10% v/v), and anaerobically using H_2_ as an energy source, and could all utilize elemental sulfur as a terminal electron acceptor. All stains also obligately required either thiosulfate or sulfur in order to utilise O_2_ as a terminal electron accepter when growing hydrogenotrophically, a trait widely observed across *Aquificota* and other hydrogenotrophic taxa (Takai, *et al*., 2001, Stohr, *et al*., 2001, Nishihara, *et al*., 1991, Kawasumi, *et al*., 1984, Dodsworth, *et al*., 2015). Sulfate reduction was not observed by any strain. Both *V. stagnispumantis* and KUI1^T^ grew in microaerophilic conditions between 0-10% O_2_ (v/v), while OKO1^T^ grew from 0-5% (v/v). Despite encoding respiratory nitrate reduction genes (NarGHI), we were unable to observe growth or a significant reduction in nitrate concentrations when tested.

### Chemotaxonomic characterizations

Respiratory quinone analysis of *V. stagnispumantis* CP.B2^T^ detected the presence of methionaquinone-7 (MTK-7). This is consistent with other *Aquificota* members, and MTK-7 was originally isolated from *Hydrogenobacter thermophilus* (Kawasumi, *et al*., 1984, Hiraishi, *et al*., 1999). The fatty acid compositions of each *Venenivibrio* strain broadly reflected that of other *Hydrogenothermaceae* taxa (Götz, *et al*., 2002, Stohr, *et al*., 2001) with profiles favouring a mixture of saturated and unsaturated C18 and C20 fatty acids and a lower concentration of short chain fatty acids. Interestingly, the fatty acid content of all three *Venenivibrio* strains were dominated by C_18:0_ and C_20:1n-9_ fatty acids, with C_18:0_ being the primary FA in OKO1^T^, whereas the opposite was true for *V. stagnispumantis* CP.B2^T^ and KUI1^T^. Four unsaturated fatty acids were detected in all strains, which is unusual for thermophilic microorganisms as they tend to favour saturated fatty acids which provide greater heat-stability for cytoplasmic membranes (Chan, *et al*., 1971, Siliakus, *et al*., 2017), however it is consistent with previous findings among *Aquificota*-specific fatty acid signatures (Jahnke, *et al*., 2001).

Catalase and oxidase tests were both negative for all strains. All isolates produced a positive Gram stain result. Despite this, transmission electron microscopy indicate a diderm membrane structure for each (Figure 1, S2), and a result consistent with all other *Aquificota* (Reysenbach, 2015). All strains encoded genes typically characteristic of Gram negative bacteria, including outer-membrane associated transporters and efflux pumps and lipopolysaccharide-associated genes (Rollauer, *et al*., 2015). Consistent with the catalase test, none of the *Venenivibrio* genomes contained genes encoding for catalase, and despite the negative oxidase test, all strains encoded genes for cytochrome C. It may be that the cytochrome C genes were not highly expressed under the conditions tested to observe a positive activity, or the pathway may have been inhibited by environmental factors (Simon and Hederstedt, 2011, Beckett, *et al*., 2000, Mendez, *et al*., 2026, Guo, *et al*., 2022). Indicator enzymatic activities were tested using the API**®** ZYM kits with *V. stagnispumantis* CP.B2^T^ was positive for napthol-AS-BI-phosphohydrolase activity, and weak acid phosphatase activity. KUI1^T^ was positive for esterase C4, esterase lipase C8, and napthol-AS-BI-phosphohydrolase, with weak acid phosphatase activity. OKO1^T^ also displayed esterase C4 and napthol-AS-BI-phosphohydrolase activity, with weak esterase lipase and acid phosphatase activity. All other enzymatic tests resulted in negative activities for the three strains.

### Taxonomic considerations

*Venenivibrio* was originally isolated in 2008 from Champagne Pool, Aotearoa-New Zealand (Hetzer, *et al*., 2008) and has since been identified as a dominant bacterial genera within Aotearoa-New Zealand geothermal systems (Power, *et al*., 2018). The wide distribution of *Venenivibrio* taxa across Aotearoa-New Zealand was inconsistent with the limited growth range of the type strain (Hetzer, *et al*., 2008). In this study, we re-characterized the type strain alongside the novel isolates to compare chemotaxonomic traits among members of the genus. We investigated the salinity, pH, and temperature tolerance ranges, as well as electron donors and acceptors, nitrogen, carbon, and sulfur sources. The results showed that the original characterisation was incomplete and that *V. stagnispumantis* CP.B2^T^ was capable of growth far beyond the ranges described. The optimum pH for the strain (pH 6.0) was greater than the reported pH_max_ in the description study (pH 5.8) and growth was observed up to pH 8.5. *V. stagnispumantis* CP.B2^T^ was reported to grow between 45-75 °C, however this study saw growth from 35-80 °C. It could also tolerate a 10 × greater concentration of NaCl (0.8% vs 8% w/v) than previously reported, and was still viable at these concentrations. Anaerobic growth using sulfur as a terminal electron acceptor was also identified, which was tested by the original authors but was not reported. In addition to these experiments, we re-sequenced the 16S rRNA gene, sequenced the whole genome, and carried out fatty acid and quinone analyses on this strain.

Morphologically, KUI1^T^ appears similar to CP.B2^T^ as curved rods, however neither KUI1^T^ or OKO1^T^ form flocs in culture as CP.B2^T^ does. OKO1^T^ displayed a predominantly swimming type motility compared to both KUI1^T^ and CP.B2^T^ which tended to spin and twitch. Genomically, KUI1^T^ is more similar to OKO1^T^ than CP.B2^T^. In addition, there are key differences in the genomes that distinguish the two new isolates from each other including the presence of a fully encoded nitrogenase (Nif operon) in OKO1^T^, and the different hydrogenase groups encoded in KUI1^T^ and OKO1^T^ (Hydrogenase group 1b v 3b respectively). The CP.B2^T^ genome possesses neither sulfur oxidation (*sox*) or nitrate reduction genes, unlike OKO1^T^ and KUI1^T^ which encode both. KUI1^T^ was capable of utilizing a broader variety of sulfur compounds as its sole sulfur source either of the other two strains (five for KUI1^T^, four for CP.B2^T^, and three for OKO1^T^).

These findings are more consistent with the widespread distribution of *Venenivibrio* taxa across geothermal environments spanning a broad range of physicochemical conditions (Power, *et al*., 2024). This physiological versatility may contribute to the ecological success of the genus and help explain its high relative abundance in many hot springs throughout the Taupō Volcanic Zone. Although comparative genomic analyses revealed substantial diversity in metabolic gene content among the strains including the fully encoded nitrogenase operon and differing hydrogenase groups, these differences were not always reflected in their phenotypes under the laboratory conditions examined. We did not observe sulfur oxidation or nitrate reduction *in vitro*. This disparity suggests that some metabolic pathways may be expressed only under specific environmental conditions or that certain genes may not be functional. Future studies examining gene expression and metabolic activity under environmentally relevant conditions will be important for determining the ecological significance of this genomic diversity and whether it contributes to the apparent endemism of *Venenivibrio* to Aotearoa-New Zealand.

## Conclusions

Each strain investigated in this study was capable of growth over a wide range of physical and chemical conditions. Genomic characterisation identified an array of metabolic genes putatively encoding sulfur oxidation, nitrogen fixation, and/or nitrate reduction across strains, although these traits were not phenotypically observed. The recharacterization of *V. stagnispumantis* CP.B2^T^ in this study differed greatly to the original characterisation, indicating a far broader tolerance of environmental conditions. This flexibility likely allows for *Venenivibrio*’s dominance in geothermal hot springs within the Taupō Volcanic Zone. Based on the differences among physiological, biochemical, and genomic traits, we propose that strains KUI1^T^ and OKO1^T^ are each novel species, and differing from the type strain, *V. stagnispumantis* CP.B2^T^. ANI values show they are closely related, but in conjunction with the phenotypic evidence provided, there is sufficient difference between them to conclude they are separate species. The substantial levels of microdiversity found among these *Venenivibrio* isolates, and previous culture-independent surveys (Power, *et al*., 2018, Power, et al., 2024) may explain *Venenivibrio* is the most abundant and prevalent bacterial genus within the Taupō Volcanic Zone.

### Emended genus description of *Venenivibrio* Hetzer et al. 2008

The description is based of Hetzer *et al.,* 2008, with the following modifications. Bacteria belonging to this genus are facultative anaerobes and are able to utilise elemental sulfur as a terminal electron acceptor in the absence of oxygen. Primary quinone MTK-7.

### Emended description of *Venenivibrio stagnispumantis* sp. nov. (Hetzer, *et al*., 2008)

*Venenivibrio stagnispumantis* (stag.ni.spu.man′tis. L. n. *stagnum* pool; L. part. adj. *spumans* foaming, frothing; N.L. gen. n. *stagnispumantis* from a frothing pool, referring to Champagne Pool).

Exhibits the following properties in addition to those described for the genus. Slightly curved rods (vibrio) with a mean length of 1.30±0.26 μm and a mean width of 0.37±0.04 μm. Grows at 38.5-80.0 °C and pH 3.0-8.5, with optimum growth at 70 °C and pH 6.0. Grows in 0-8.0% (w/v) NaCl; optimum growth occurs at 0.8% (w/v) NaCl. Oxygen is the preferred terminal electron acceptor, but can utilize sulfur in the absence of oxygen. Tolerates oxygen in the range 1-10 % (v/v). Multiple flagella present. Fatty acid composition: C_20:1n-9_ (46.9%), C_18:0_ (31.1%), C_18:1n-9_ (10.5%), C_20:0_ (4.2%), C_12:0_ (2.9%), C_16:0_ (1.6%), C_18:1n-7_ (1.5%), C_20:1n-7_ (1.1%). Primary quinone: methionaquinone-7. Genome size 1.56 Mbp,

The type strain is CP.B2^T^ (=JCM 14244^T^ =DSM 18763^T^), isolated from the terrestrial hot spring Champagne Pool in Waiotapu, New Zealand. The DNA G+C content of the type strain is 29.2 mol%.

## Declaration of competing interests

The authors declare no competing financial or personal interests that could influence this work reported in this paper.

## Acknowledgments

The authors would like to thank Ngāti Tahu-Ngāti Whaoa for their support of this research and access to sampling locations. Ngāti Tahu-Ngāti Whaoa have mana whenua (customary rights) over Orakei Korako and Waiotapu, where *Venenivibrio orakeikorakoensis* and *Venenivibrio stagnispumantis* CP.B2^T^ were isolated, and all data associated with these strains. We thank them for gifting the name for *V. orakeikorakoensis*. The authors also thank the Rotorua District Council for permission to sample Kuirau Park. We also wish to thank Marina Richena, Kim Parker, and Duane Harland for their assistance with transmission electron microscopy performed at AgResearch, New Zealand.

## Notes

### Competing Interest Statement

The authors have declared no competing interest.

### Funding

- New Zealand Royal Society Marsden grant: MFP-UOC2201

