## Supplementary Data for "Description of *Venenivibrio orakeikorakoensis* sp. nov. and *Venenivibrio aotearoaensis* sp. nov, and emended description of genus *Venenivibrio*, *Venenivibrio stagnispumantis* species"

Supplementary Table 1: BLAST identity pairwise matrix of 16S rRNA gene sequences between the two novel *Venenivibrio* species, *V. stagnispumantis,* and representative members of each genus within *Hydrogenothermaceae.* Values are percentages. Accession numbers for reference genes are *V. stagnispumantis* CP.B2^T^ (NR_044029.1)*, Sulfurihydrogenibium azorense* Az-Fu 1^T^ (NR_102858.1)*, Persephonella hydrogeniphila* 29W^T^ (NR_024797.1), and *Hydrogenothermus marinus* VM1^T^ (NR_114754.1).

|  | OKO1 | KUI1 | *V. stagnispumantis* CP.B2^T^ | *S. azorense* Az-Fu1^T^ | *P. hydrogeniphila* 29W^T^ | *H. marinus* VM1^T^ |
| --- | --- | --- | --- | --- | --- | --- |
| OKO1 | - | 99.48 | 99.13 | 94.65 | 90.96 | 89.09 |
| KUI1 | 99.48 | - | 98.82 | 95.13 | 90.95 | 89.13 |
| *V. stagnispumantis* CP.B2^T^ | 99.13 | 98.82 | - | 94.45 | 89.91 | 88.96 |
| *S. azorense* Az-Fu1^T^ | 94.65 | 95.13 | 94.45 | - | 90.93 | 89.72 |
| *P. hydrogeniphila* 29W^T^ | 90.96 | 90.96 | 89.91 | 90.93 | - | 92.38 |
| *H. marinus* VM1^T^ | 89.09 | 89.13 | 88.96 | 89.72 | 92.38 | - |

Supplementary Table 2: Whole genome analyses of the two novel *Venenivibrio* species*, V. stagnispumantis,* and representative members of each genus within *Hydrogenothermaceae.* Accession numbers for reference strains are *V. stagnispumantis* CP.B2^T^*, Sulfurihydrogenibium azorense* Az-Fu 1^T^ (GCF_000021545)*, Persephonella hydrogeniphila* 29W^T^ (GCF_900215515), *Hydrogenothermus marinus* VM1^T^ (GCF_003688665). “N.D.“ indicates absent in genome. Hydrogenases classified using HydDB (Søndergaard*, et al.*, 2016)

|  | CP.B2^T^ | OKO1^T^ | KUI1^T^ | *S. azorense* Az-Fu1^T^ | *P. hydrogeniphila* 29W^T^ | *H. marinus* VM1^T^ |
| --- | --- | --- | --- | --- | --- | --- |
| Genome characteristics | | | | | | |
| Genome size | 1,566,079 | 1,735,438 | 1,690,398 | 1,640,877 | 1,998,302 | 1,604,000 |
| G+C% | 29.15 | 29.15 | 29.2 | 32.75 | 35.11 | 29.68 |
| Contigs | 89 | 55 | 24 | 1 | 19 | 18 |
| Genome quality | | | | | | |
| N50 | 65682 | 120414 | 223176 | 1640877 | 151898 | 372577 |
| N75 | 35128 | 77285 | 159365 | 1640877 | 100747 | 174834 |
| L50 | 9 | 6 | 3 | 1 | 3 | 2 |
| L75 | 17 | 10 | 5 | 1 | 7 | 4 |
| Completeness (%) | 99.59 | 99.59 | 99.59 | 99.39 | 99.39 | 98.98 |
| Contamination (%) | 1.63 | 0.41 | 0.81 | 0.0 | 0.81 | 1.73 |
| Key metabolic genes and enzyme groups | | | | | | |
| Hydrogenase groups | NiFe 2d | NiFe 2d, 3b | NiFe 1b, 2d | NiFe 2d, 3b | NiFe 1b, 2d, 3b | 1b, 2d |
| Cytochrome groups | C, bd, d | C, d | C, bd, d | C, d | C, d | C, d |
| Sulfur oxidation genes | N.D. | soxABXYZ | soxABXYZ | soxABXYZ | soxABXYZ | soxABXYZ |
| Respiratory Nitrate reduction genes | N.D. | NarGHI | NarGHI | NarH | NarGH | NarGH |
| Nitrogen fixation | N.D. | NifABDEHKNXU | N.D. | N.D. | N.D. | N.D. |

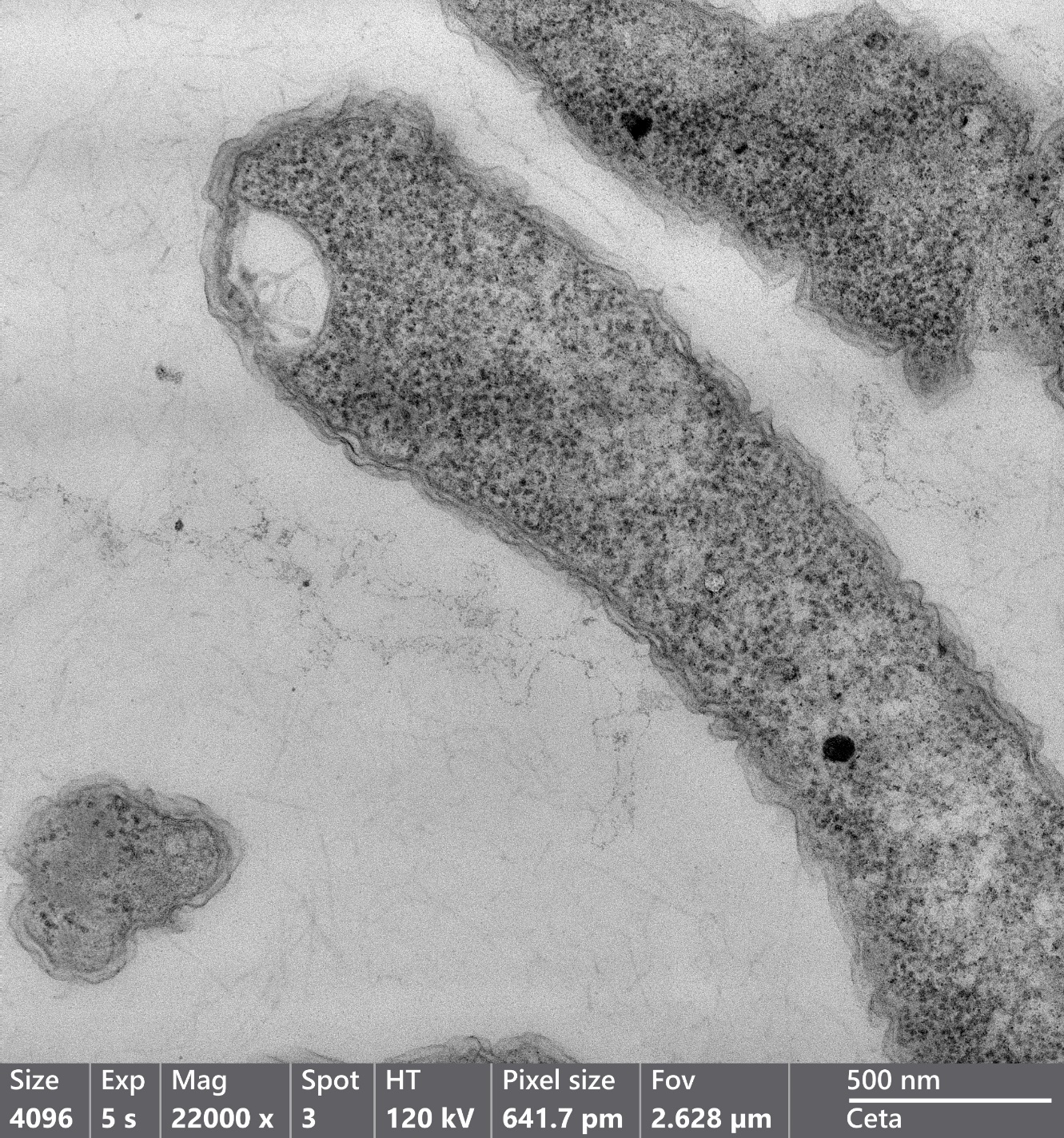

Supplementary Figure 1: Thin section of V. stagnispumantis CP.B2^T^ viewed at 22,000 × magnification (1% uranyl acetate, 30 sec exposure time). Scale bar indicates scale of 500 nm.

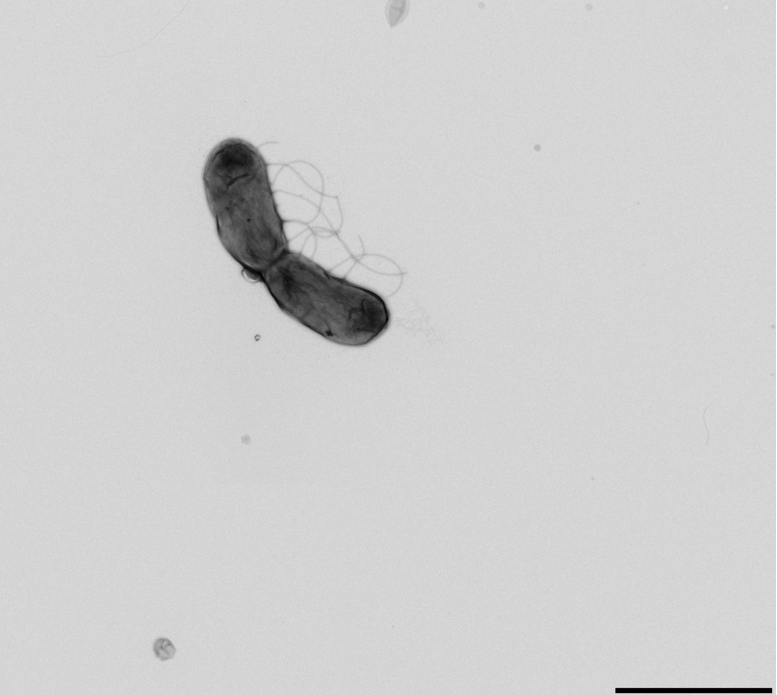

Supplementary Figure 2: Transmission electron micrograph of negative stained Venenivibrio aotearoaensis KUI1^T^ (1% uranyl acetate, 30 sec exposure time) at 4300 × magnification. Scale bar represents scale of 2 μm.

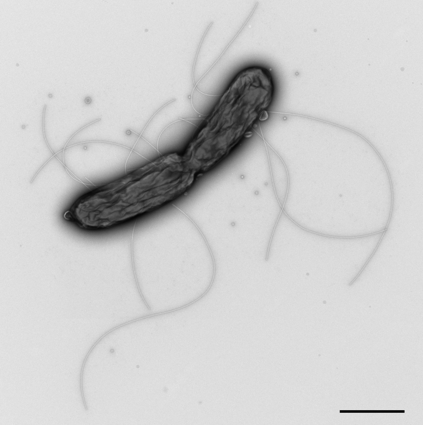

Supplementary Figure 3: Transmission electron microscopy image of negative stained V. stagnispumantis CP.B2^T^ (0.5% uranyl acetate, 10 sec exposure time) at 8500 × magnification. Scale bar represents scale of 1 μm.

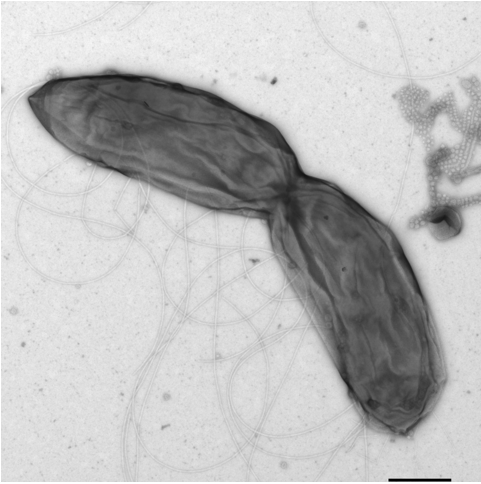

Supplementary Figure 2: Transmission electron microscopy image of negative stained Venenivibrio orakeikorakoensis OKO1^T^ (0.5 % w/v uranyl acetate, 10 sec exposure time) at 13,500 × magnification. Scale bar represents scale of 500 nm.

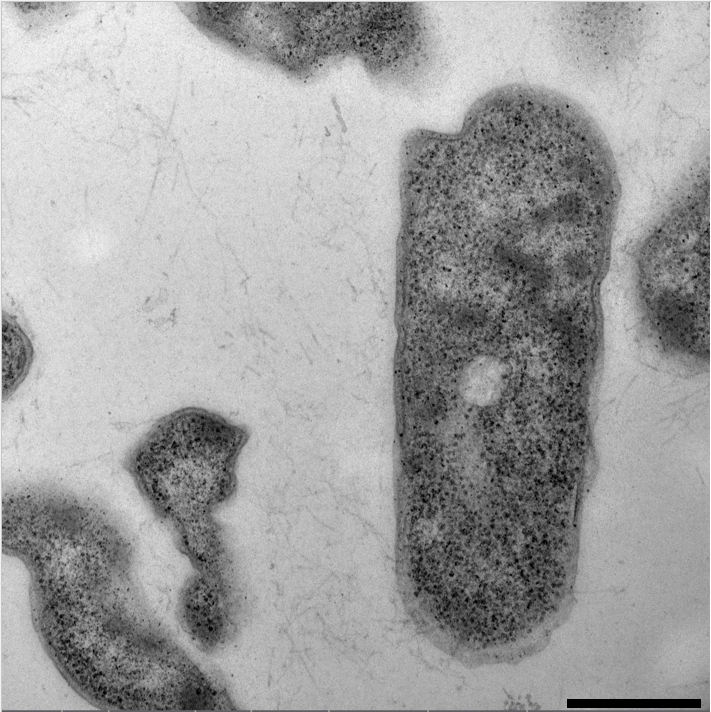

Supplementary Figure 3: Transmission electron micrograph of Venenivibrio orakeikorakoensis OKO1^T^ thin section (0.5% w/v uranyl acetate, 10 sec exposure time) at 22,000 × magnification. Scale bar represents scale of 500 nm.
